# Reconciling Base-Pair Resolution 5-methylcytosine and 5-hydroxymethylcytosine Data in Neuroepigenetics

**DOI:** 10.1101/630772

**Authors:** Joseph Kochmanski, Candace Savonen, Alison I. Bernstein

**Affiliations:** Department of Translational Science and Molecular Medicine, College of Human Medicine, Michigan State University, Grand Rapids, MI, United States

## Abstract

Epigenetic marks operate at multiple chromosomal levels to regulate gene expression, from direct covalent modification of DNA to 3D chromosomal structure. Research has shown that 5-methylcytosine (5-mC) and its oxidized form, 5-hydroxymethylcytosine (5-hmC), are stable epigenetic marks with distinct genomic distributions and separate regulatory functions. In addition, recent data indicate that 5-hmC plays a critical regulatory role in the mammalian brain, emphasizing the importance of considering this alternative DNA modification in the context of neuroepigenetics. Traditional bisulfite (BS) treatment-based methods to measure the methylome are not able to distinguish between 5-mC and 5-hmC, meaning much of the existing literature does not differentiate these two DNA modifications. Recently developed methods, including Tet-assisted bisulfite (TAB) treatment and oxidative bisulfite (oxBS) treatment, allow for differentiation of 5-hmC and/or 5-mC levels at base-pair resolution when combined with next-generation sequencing or methylation arrays. Despite these technological advances, there remains a lack of clarity regarding the appropriate statistical methods for integration of 5-mC and 5-hmC data. As a result, it can be difficult to determine the effects of an experimental treatment on 5-mC and 5-hmC dynamics. Here, we propose a statistical approach involving mixed effects to simultaneously model paired 5-mC and 5-hmC data as repeated measures. Using this approach, it will be possible to determine the effects of an experimental treatment on both 5-mC and 5-hmC at the base-pair level.

## 1. Introduction

### 1.1 Epigenetics

Epigenetic marks operate at four major levels – DNA modifications, histone modifications, non-coding RNAs, and 3-D chromatin structure (Chen et al., 2017b). The most studied DNA modification is 5-methylcytosine (5-mC), the addition of a methyl group at the C5 position of a cytosine in the DNA sequence (Moore et al., 2013). An abundance of research shows associations between 5-mC and gene expression, and suggests that this epigenetic mark plays a key role in transcriptional control (Moore et al., 2013). In addition to 5-mC, there are three further oxidized DNA modifications – 5-hydroxymethylcytosine (5-hmC), 5-formylcytosine (5-fC), and 5-carboxylcytosine (5-caC) (Shen et al., 2014). These alternative DNA modifications are formed when 5-mC is successively oxidized by the ten-eleven translocase (Tet) family of proteins (Shen et al., 2014). The 5-fC and 5-caC modifications are rapidly removed by thymine-DNA glycosylase (TDG) and base excision repair, and are thought to be transient (He et al., 2011; Ito et al., 2011; Maiti and Drohat, 2011). In contrast, 5-hmC can be a stable epigenetic mark that regulates transcription (Hahn et al., 2014). In particular, 5-hmC appears to play an important role in the central nervous system, where it is present at much higher levels than embryonic stem cells and other somatic tissues (Cheng et al., 2015; Globisch et al., 2010; Nestor et al., 2012; Szwagierczak et al., 2010).

### 1.2 Neuroepigenetics: a unique role for 5-hmC

Given the relative enrichment of 5-hmC in nervous tissue, an abundance of new research has examined the potential regulatory role of 5-hmC in the brain. Studies show that 5-hmC is acquired during neuronal development (Hahn et al., 2013; Szulwach et al., 2011) and maintained throughout adulthood (Chen et al., 2014). In the brain, 5-hmC has a specific distribution across the genome, with enrichment at genic regions, distal regulatory elements, and exon-intron boundaries (Khare et al., 2012; Lister et al., 2013; Wen et al., 2014). At the level of specific genes, 5-hmC is enriched in gene bodies of genes that are transcriptionally active in neuronal tissue (Mellén et al., 2012). In addition, different anatomical regions of the brain show distinct 5-hmC patterning (Lunnon et al., 2016), suggesting a specific regulatory role for this epigenetic mark.

Recent work also highlights that 5-mC and 5-hmC differ in their genomic distribution in the nervous system (Chen et al., 2014; Cheng et al., 2015). During synaptogenesis, 5-hmC preferentially accumulates in euchromatin, whereas 5-mC gradually builds up in heterochromatic regions (Chen et al., 2014). In addition, 5-mC and 5-hmC preferentially recruit distinct sets of DNA binding proteins in brain tissue (Spruijt et al., 2013). For example, whereas Mbd1, Mbd4, and MeCP2 bind 5-mC at higher affinity, Neil1, Thy28, and Wdr76 have a higher affinity for 5-hmC (Spruijt et al., 2013). 5-hmC is also preferentially bound by the DNA-binding protein Uhrf2 in neuronal progenitor cells (Spruijt et al., 2013), a process that may regulate spatial memory and learning (Chen et al., 2017a). The distinct sets of readers for 5-hmC and 5-mC indicate that these two epigenetic marks have separate regulatory functions in neuronal tissue.

Combined, the available data suggest that 5-hmC plays a critical regulatory role in the mammalian brain, emphasizing the importance of considering this alternative DNA modification in the context of neuroepigenetics. As such, it is critical that the field develops methods to accurately distinguish 5-hmC from 5-mC in a genome-wide context. Here, we discuss the available methods for measuring 5-hmC, including their strengths and weaknesses, and then propose a statistical approach for co-analyzing the effects of an experimental treatment on paired 5-mC and 5-hmC data.

## 2. Base-pair resolution differentiation of 5-mC and 5-hmC

Historically, the majority of neuroepigenetics studies investigating DNA modifications utilized bisulfite (BS) treatment-based methods to measure DNA methylation (Beck and Rakyan, 2008; Clark et al., 2006; Rein et al., 1998). BS conversion utilizes sodium bisulfite to convert all unmodified cytosines to uracil by deamination, but does not deaminate 5-mC or 5-hmC. The converted cytosines (C, 5-fC or 5-caC) are read as thymines during sequencing, while the unconverted cytosines (5-mC or 5-hmC) are read as cytosines. From this data, the percent of methylation (beta value) at each cytosine can be calculated from the proportion of cytosines and thymines detected at each position (Figure 1).

**Figure 1.**
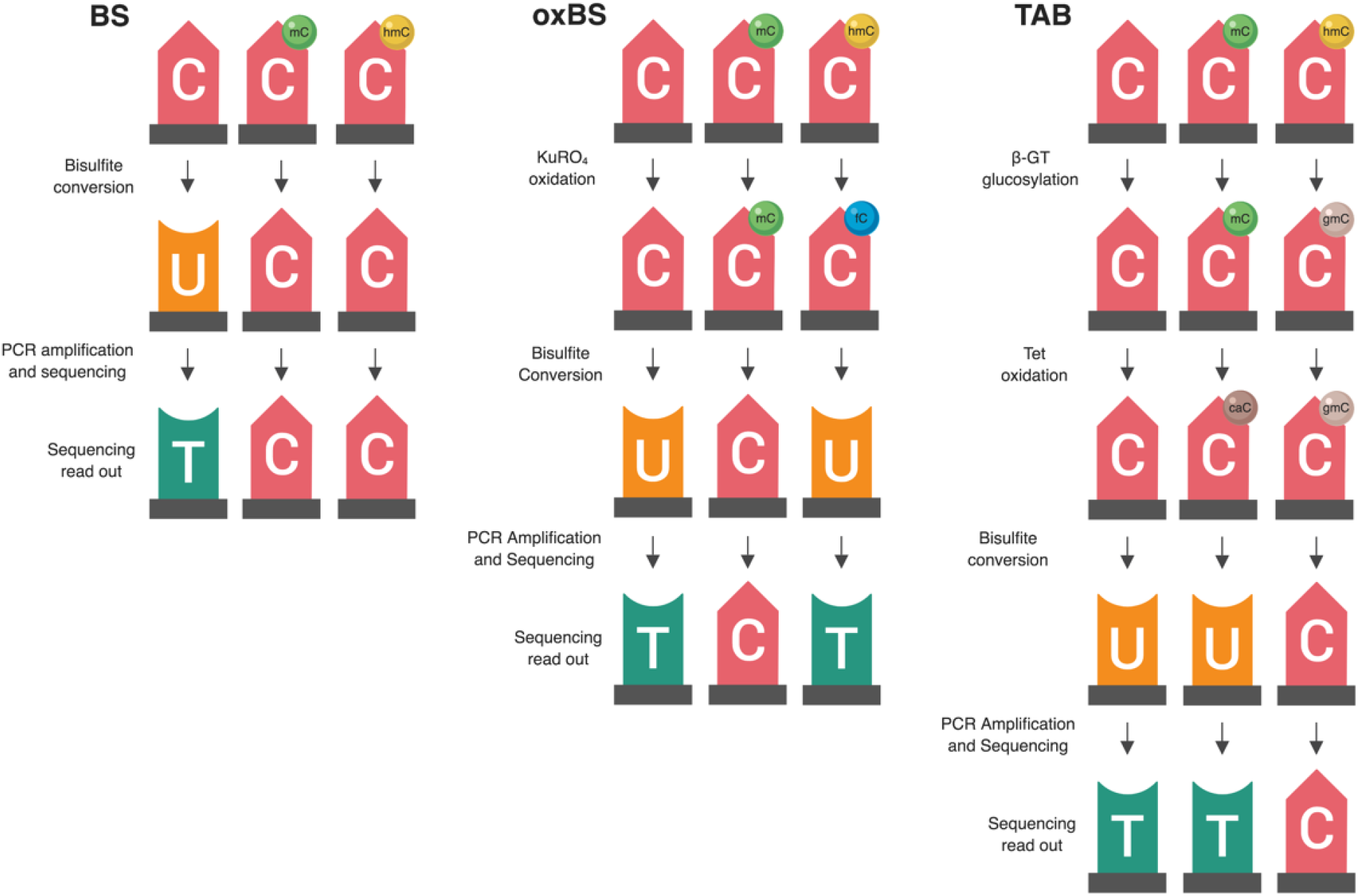
Summary of available methods for measuring genomic 5-hmC levels. There are two widely-adopted methods used to measure 5-hmC levels at the base pair level – paired bisulfite/oxidative bisulfite treatment (BS/oxBS) and TET-assisted bisulfite treatment (TAB) (Booth et al., 2013; Yu et al., 2012). These three methods differ in their chemistry and data interpretation. In the oxBS method, KRuO_4_ oxidation selectively converts 5-hmC to 5-fC, which is removed during bisulfite conversion. By comparing oxBS data (5-mC) with traditional BS data, it is possible to infer 5-hmC levels. For the TAB treatment method, 5-hmC is selectively tagged with a β-glucosyl group, which makes it resistant to either bisulfite conversion. One its own, TAB provides a true value for 5-hmC, but does not measure 5-mC.

Unfortunately, 5-mC and 5-hmC are both resistant to deamination during BS conversion, meaning BS-based methods are unable to differentiate between these two marks (Huang et al., 2010; Jin et al., 2010). As such, studies utilizing traditional BS treatment actually captured both 5-mC and 5-hmC, which may confound their identified associations between differential DNA methylation and transcriptional control. To address this issue, multiple technological advancements have allowed for specific profiling of 5-mC and 5-hmC at the base pair level. Currently, there are two BS treatment-based methods used to measure 5-hmC levels – oxidative bisulfite treatment (oxBS) and TET-assisted bisulfite treatment (TAB). More recently, additional techniques have been developed to estimate true 5-mC and true 5-hmC values, including APOBEC3A-mediated deamination sequencing, TET-assisted pyridine borane sequencing, and AbaSI-sequencing (Li et al., 2018; Liu et al., 2019; Sun et al., 2013). These alternate methods hold promise, but have not yet been widely adopted by the field. While this article focuses on a new statistical approach to deal with paired 5-mC and 5-hmC data from bisulfite treatment-based methods, the proposed approach should apply to any set of methods that generates paired 5-mC and 5-hmC data.

### 2.1 Bisulfite/oxidative bisulfite treatment (BS/oxBS)

Oxidative bisulfite (oxBS) treatment involves chemically-mediated selective oxidation of 5-hmC to 5-fC prior to BS conversion by potassium perruthenate (KuRO_4_). After this oxidation step, 5-hmC acts like 5-fC during BS conversion and is converted to uracil and read as thymine in subsequent sequencing reactions. 5mC remains unaffected by KuRO_4_, is not deaminated by BS and is read as cytosine. Thus, oxBS provides a measure of 5-mC only (“true 5-mC”) (Booth et al., 2013) (Table 1). This method must be paired with traditional BS conversion, which provides a combined measure of 5-mC and 5-hmC, and an estimation step must be performed to generate an estimate of 5hmC. Currently, paired BS/oxBS is the most commonly used method to generate paired 5-mC and 5-hmC data; it is standard practice to use a maximum likelihood estimate method to estimate 5hmC levels from paired BS/oxBS data (Xu et al., 2016).

**Table 1.**
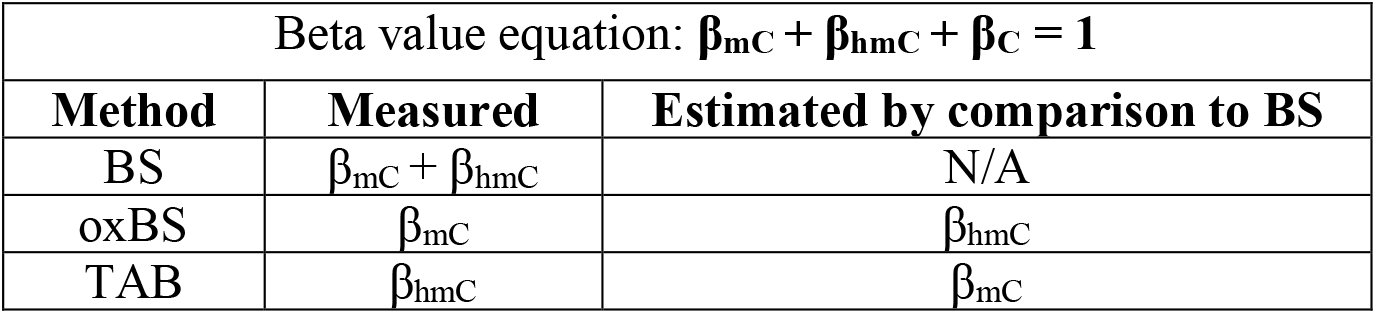
Beta value estimation for each described method for measuring base-pair resolution 5-hmC. The various DNA treatment methods described in the text – BS, oxBS, and TAB – allow for specific tagging and measurement of different DNA modifications. By comparing beta values generated from these methods to bisulfite treatment data, β_hmC_ and β_mC_ can be estimated. As indicated by the equation at the top of the table, the sum of beta values for all DNA modifications is always equal to 1. This is because beta values represent the proportions of each modification, not measures of magnitude. As a result of being proportions, β_hmC_ and β_mC_ will always have values between 0-1.

| Beta value equation: $\beta_{mC} + \beta_{hmC} + \beta_C = 1$ | | |
| --- | --- | --- |
| Method | Measured | Estimated by comparison to BS |
| BS | $\beta_{mC} + \beta_{hmC}$ | N/A |
| oxBS | $\beta_{mC}$ | $\beta_{hmC}$ |
| TAB | $\beta_{hmC}$ | $\beta_{mC}$ |

### 2.2 TET-assisted bisulfite treatment (TAB)

TET-assisted bisulfite treatment (TAB) method is an enzyme-based method where 5-hmC is specifically protected from ten-eleven translocase (TET) enzyme-mediated oxidation. In this method, a β-glucosyltransferase enzyme is used to add a glucose moiety to 5-hmC prior to treatment with recombinant TET enzyme. The TET enzyme oxidizes 5-mC, but not glucosylated 5-hmC, to 5-caC, a DNA modification that can be bisulfite converted (Yu et al., 2012).

In essence, this method selectively protects 5-mC, leaving 5-hmC and the other modified cytosines available for BS conversion. As a result, TAB directly measures 5-hmC (“true 5-hmC”) at the base-pair level and can be performed without paired BS conversion (Table 1). However, the TAB method only measures 5-hmC, and does not provide any information on 5-mC. In addition, the enzymatic treatment required for TAB can be quite costly. Based on these considerations, use of the TAB method remains limited compared to BS/oxBS treatment.

### 2.3 Generation of 5-mC and 5-hmC beta values

For genome-wide assessment of 5-mC and 5-hmC, each of the methods described above can be paired with sequencing arrays (i.e. Illumina 450K/EPIC BeadChip), reduced representation sequencing, or whole-genome sequencing. Choosing between these available methods is not only a question of cost, but also experimental question, tissue type, and desired genomic coverage. Discussion of these specific issues is beyond the scope of this commentary; they are discussed in depth elsewhere (Kurdyukov and Bullock, 2016; Sun et al., 2015; Yong et al., 2016). The issues related to co-analysis of 5-mC and 5-hmC exist for all three types of data generation.

Following conversion of DNA by any of these three methods and subsequent analysis by sequencing arrays (i.e. Illumina 450K/EPIC BeadChip), reduced representation sequencing, or whole-genome sequencing, beta values for each modification can be calculated at each assayed cytosine. Beta values are ratios of modified (5-mC or 5-hmC) and unmodified (C) alleles, with values between 0 (unmodified) and 1 (fully modified); added together, the sum of these beta values at each cytosine equals 1 (Table 1).

## 3. Problems in 5-mC and 5-hmC data analysis

Despite the significant technological advances in differentiating 5-mC and 5-hmC, standard statistical methods for co-analyzing 5mC and 5hmC do not yet exist. At an individual CpG site, both 5-mC and 5-hmC can contribute to gene regulation, but none of the available bioinformatics tools provide a function for co-analyzing 5-mC and 5-hmC β values. As a result, existing studies have focused on either examining the distribution of 5-hmC across the genome in isolation (Green et al., 2016; Hernandez Mora et al., 2018; Johnson et al., 2016) or treating 5-mC and 5-hmC β values as independent variables, analyzing each epigenetic mark as a separate dataset to identify differentially methylated and hydroxymethylated regions (Glowacka et al., 2018; Zhang et al., 2018). While there is utility to both of these approaches, the results are difficult to reconcile into a clear picture of the underlying biology for two main reasons: 1) the methodological and biological interdependence of 5-mC and 5-hmC and 2) the different distributions of β_mC_ and β_hmC_. This uncertainty complicates functional interpretation of BS-based DNA modification data, since 5-mC and 5-hmC have distinct genomic distributions and regulatory functions (Shen and Zhang, 2013; Skvortsova et al., 2017). Furthermore, this type of differential DNA modification misclassification is particularly relevant in nervous system tissue, where 5-hmC is present at high levels (Cheng et al., 2015; Globisch et al., 2010; Nestor et al., 2012; Szwagierczak et al., 2010). Below, we run through these concerns in greater detail, and then propose a statistical method for co-analyzing paired 5-mC and 5-hmC levels.

### 3.1 Interdependence of 5-mC and 5-hmC

After measuring genome-wide 5-mC and 5-hmC at the base-pair level, a simple approach would be to split these two epigenetic marks into separate datasets for analysis. While this method is attractive, it fails to account for the interdependence of 5-mC and 5-hmC data. These two epigenetic marks are often related to each other biologically and methodologically. Biologically, 5-hmC is produced through direct oxidation of 5-mC (Shen et al., 2014), meaning 5-hmC β values are directly dependent on 5-mC β values. In addition to their biological relationship, 5-mC and 5-hmC β values generated from BS/oxBS experiments are also methodologically related, since calculation of 5-hmC is dependent upon either subtraction or a maximum likelihood estimation step (Booth et al., 2013; Houseman et al., 2016; Xu et al., 2016). Unless one were to measure 5-mC and 5-hmC directly through an alternative combination of the presented techniques, this methodological interdependence is unavoidable. Modeling approaches that treat 5-mC and 5-hmC β values as independent variables do not account for this inherent interdependence, and limit one’s ability to comprehensively identify regions where 5-mC and 5-hmC have differential responses to an experimental condition.

### 3.2 Differential distributions of β_mC_ and β_hmC_

Even in the brain, where 5-hmC is present at comparatively high levels, it is still a rare event. Thus, many CpG sites have appreciable 5-mC, but no 5-hmC, which means that estimated 5-hmC β values are zero-enriched (Figure 2). On a genome-wide scale, 5-mC has a beta distribution and 5-hmC has a zero-inflated beta distribution. Given these divergent distributions, independent tests for differential 5-mC and 5-hmC need to utilize specific statistical approaches that include appropriate assumptions for their distributions. When different statistical tests are used for 5-mC and 5-hmC, the results from differential testing are difficult to reconcile. Furthermore, zero values for 5-hmC are typically estimated from paired BS/oxBS data, so it can be difficult to determine whether 5-hmC β values are true biological zeroes or technical artifacts of the data generation method. This complicates the downstream identification of treatment-induced active demethylation at specific genomic regions.

**Figure 2.**
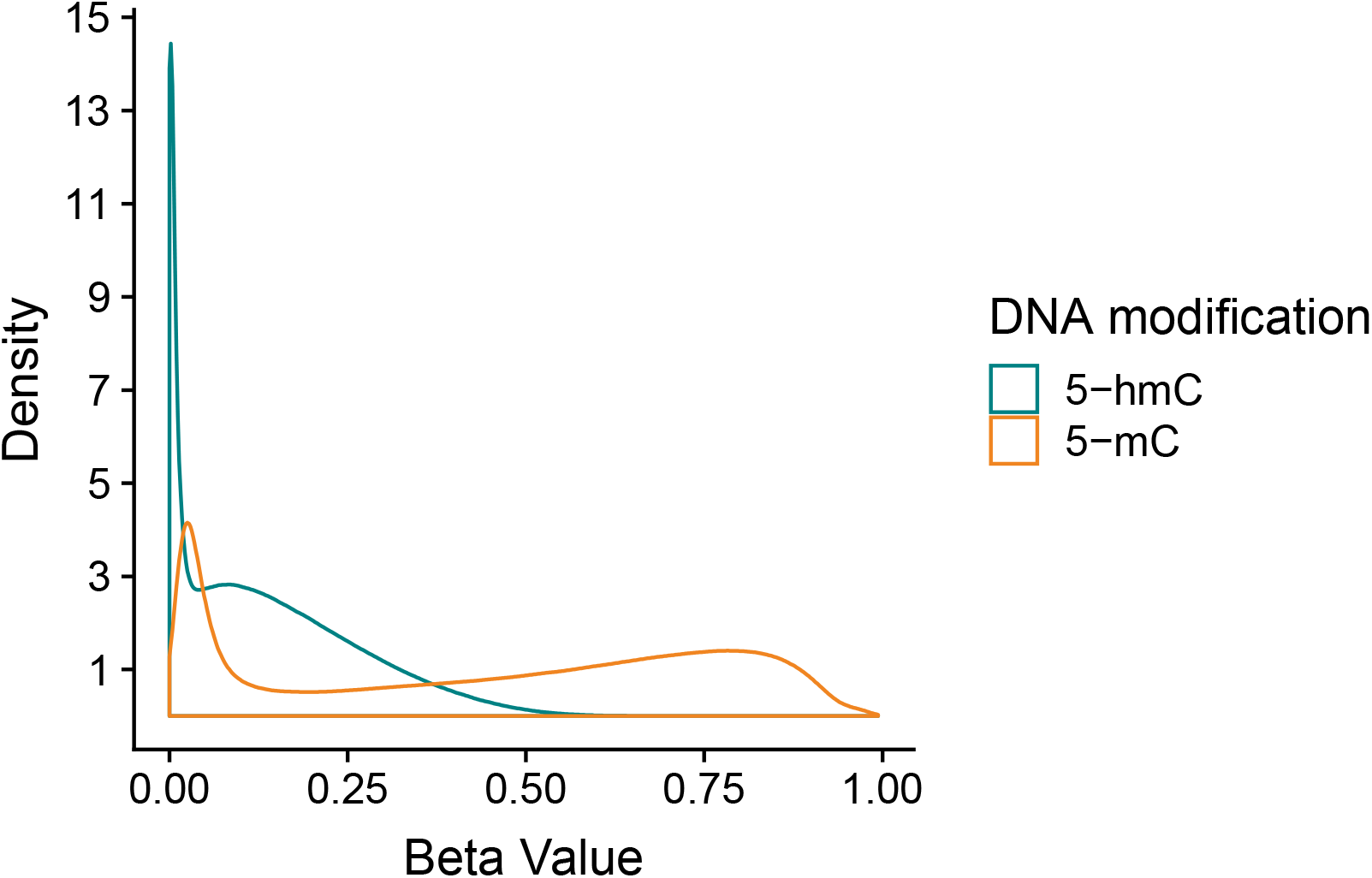
Beta value distributions for 5-mC and 5-hmC from example BS-EPIC/oxBS-EPIC array data. Beta values for 5-mC and 5-hmC were estimated from in-house example BS/oxBS-EPIC data using the oxBS.MLE function with default parameters in the *ENmix* R package. Since many CpG sites have appreciable 5-mC, but no 5-hmC, estimated 5-hmC beta values are zero-enriched after maximum likelihood estimation from BS/oxBS data.

### 3.3 Scenarios where independent analysis breaks down

Modeling 5-mC and 5-hmC data separately requires a larger number of statistical tests than analyzing a single dataset. This increases the risk for false positives, and may impede accurate interpretation of the data. While multiple testing correction methods can be used to address this concern, these statistical techniques can drastically limit one’s ability to detect true positives, especially in studies with a small sample size. As a result, analyzing 5-mC and 5-hmC data using separate models could negatively impact the ability of a project to identify regions of differential methylation and hydroxymethylation.

In addition to potential statistical errors, there are multiple scenarios in which independent analysis of 5-mC and/or 5-hmC could fail to capture a complete picture of differential DNA modifications (Figure 3). Here, we present two potential scenarios in which independent analysis of 5-mC and 5-hmC presents limitations to the biological interpretation of the results.

**Figure 3.**
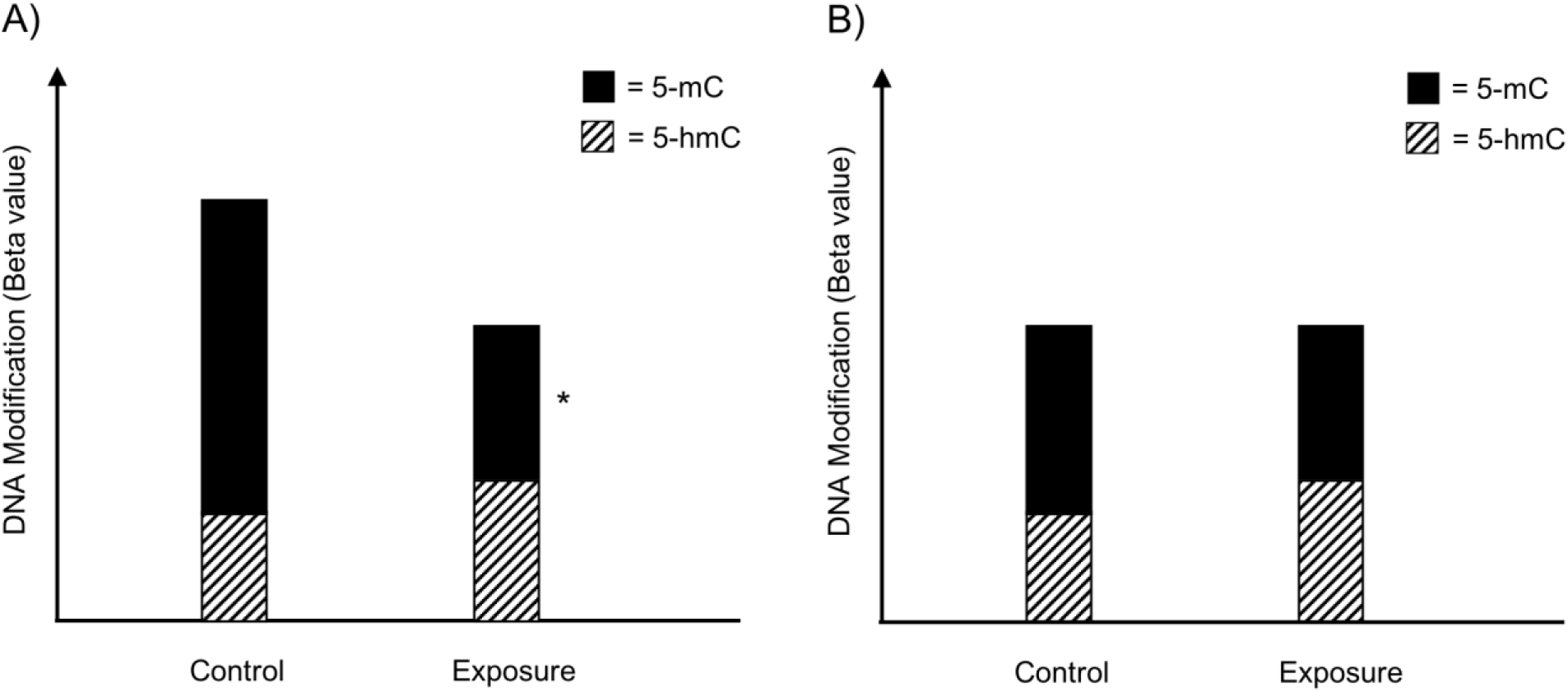
Examples of simultaneous, treatment-related changes in 5-mC and 5-hmC. Here, we present two hypothetical scenarios in which experimental condition alters levels of 5-mC and/or 5-hmC in brain tissue. A) In the first example, 5-mC significantly decreases and 5-hmC shows a non-significant increase at a CpG site, while combined levels of DNA modifications decrease in exposed compared to control. B) In the second example, 5-mC shows a non-significant decrease and 5-hmC shows a non-significant increase at a CpG site; meanwhile, combined DNA modifications remain the same by experimental group. Asterisk indicates significant changes by experimental condition.

In a first hypothetical scenario, the total proportion of modified cytosines decreases at a given CpG site, but only one modification is identified as statistically significant, leading to an incomplete view of the underlying biology (Figure 3A). In the specific example provided, the proportion of 5-mC significant decreases and the proportion of 5-hmC shows a non-significant increase. These example data suggest oxidation of 5-mC to 5-hmC at the measured CpG site. This oxidative processing may be part of active demethylation, which would lead to the observed decrease in total DNA modifications. However, downstream statistical analysis that treats 5-mC and 5-hmC as independent measures would only pick up the significant changes in 5-mC, and would likely not identify the corresponding directional shift in 5-hmC. As a result, the selected analysis approach could lead to improper biological interpretation of the results.

In a second scenario, 5-mC shows a non-significant decrease and 5-hmC shows a non-significant increase; meanwhile, combined DNA modifications remain the same by experimental group (Figure 3B). These data suggest a region with subtle oxidative processing of 5-mC to 5-hmC, but this shift in DNA modifications would not be detected in downstream statistical analysis that treats 5-mC and 5-hmC as independent measures.

For the described hypothetical scenarios, changes in the balance between 5-mC and 5-hmC at a measured CpG site may not be detected if the individual DNA modifications were analyzed as independent datasets. These dynamic regions of active DNA modification cycling may play an important biological role, and should not be ignored. To address these concerns, researchers need a method to simultaneously analyze 5-mC and 5-hmC levels; unfortunately, no such statistical method currently exists in the literature.

## 4. Potential solutions

### 4.1 Measuring true 5-mC and 5-hmC

One way to address some of the statistical concerns brought up in the previous section would be to measure true levels of 5-mC and 5-hmC. Direct measurement of 5-mC and 5-hmC would bypass the required estimation step, thereby reducing the methodological interdependence of 5-mC and 5-hmC. However, this type of combined approach does not address the statistical concerns laid out above. Furthermore, current methods are reliant upon bisulfite conversion, which negatively impacts DNA quality. This loss of sample integrity complicates integration of data generated from these BS-based methods, due to inconsistent genomic coverage. Ideally, further work in the field will lead to development and adoption of reliable methods to measure 5-mC and 5-hmC directly and independently without a harsh bisulfite conversion step to allow for consistent genomic coverage.

### 4.2 Statistical methods to analyze 5-mC and 5-hmC as related measures

Here, we propose a new approach for modeling paired 5-mC and 5-hmC data (Figure 4). Rather than treating β_mC_ and β_hmC_ as independent variables, we propose treating these two data points as “repeated” measures of a single outcome variable – “DNA modification.” To achieve this in statistical terms, we propose a mixed effects (ME) modeling approach (Laird and Ware, 1982). Under this approach, each model would include a fixed effect for experimental condition/group and random effects for CpG probe ID and batch to account for within-site and within-batch variability. Given that only two data points (5-mC and 5-hmC beta value) are included in each model per sample, inclusion of a random effect for CpG probe ID also accounts for within-sample variability. To determine the differential effects of an experimental condition on 5-mC and 5-hmC, an interaction term between experimental condition and a categorical DNA modification variable (DNA_mod_cat: “5-mC” or “5-hmC”) would also be included in the ME model. This interaction term determines whether the direction of the relationship between β values and experimental condition varies by DNA modification category (5-mC/5-hmC).

**Figure 4.**
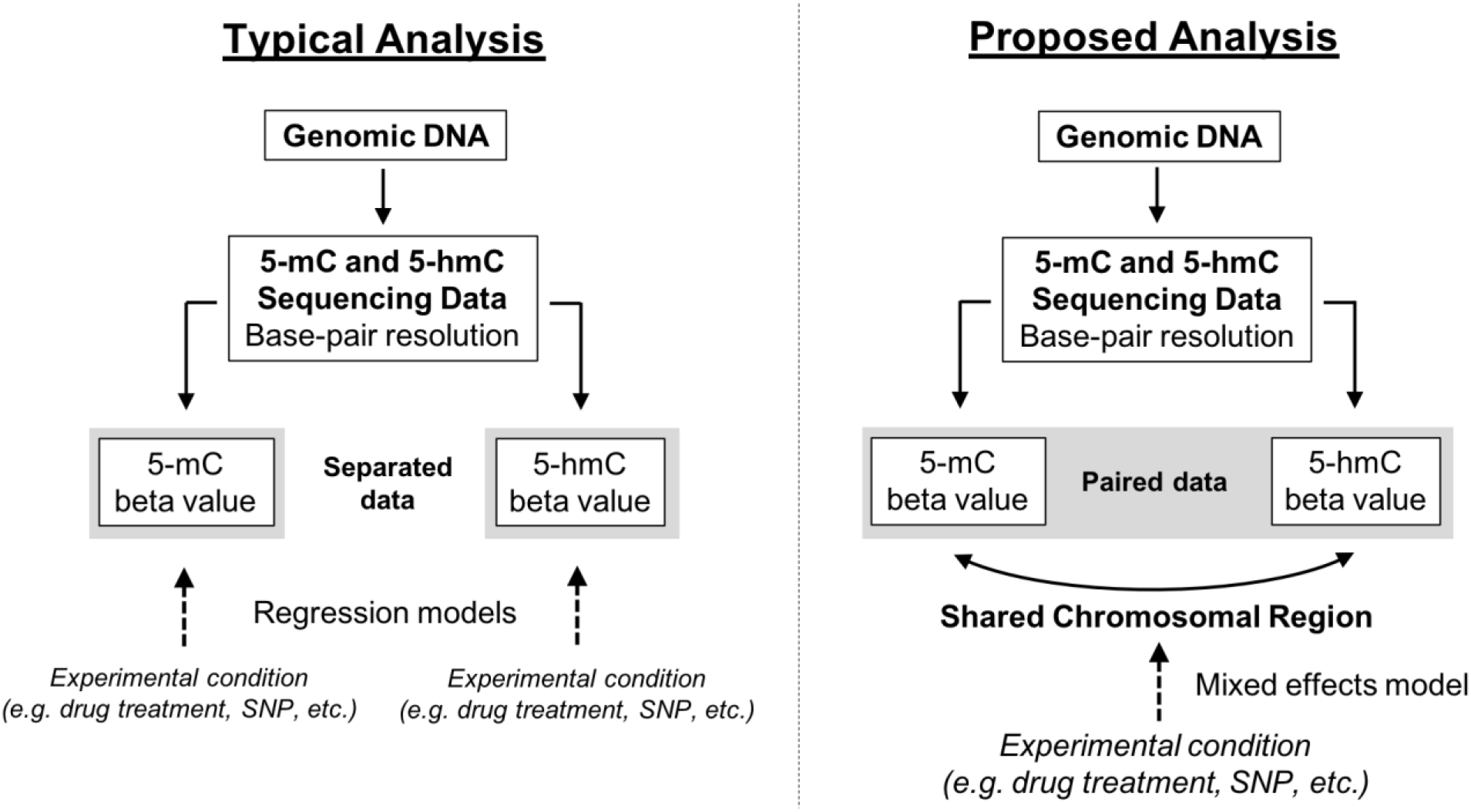
Conceptual framework for reconciling 5-mC and 5-hmC data using a mixed effects modeling approach. Upon generating base-pair resolution 5-mC and 5-hmC data, the current analysis approach is to run separate regression models on the beta values for 5-mC and 5-hmC. However, this approach fails to account for the inherent dependence of 5-hmC on 5-mC, and limits a researcher’s ability to quickly and comprehensively identify regions where 5-mC and 5-hmC have differential responses to an experimental condition. Here, we propose an alternative mixed effects modeling approach in which the 5-mC and 5-hmC beta values are treated as repeated measures. Using an interaction term in our proposed model, it will be possible to identify regions where the directionality of response to experimental condition changes by DNA modification (5-mC vs. 5-hmC). This approach is not necessarily a replacement for separate regression models, but rather a supplement.

Mixed models could be fit using a model in the following form:

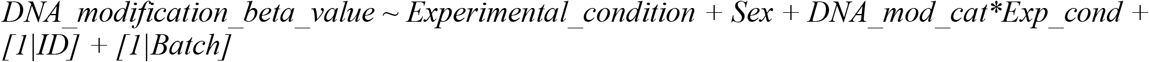

Alternatively, the main effect of experimental condition on DNA modifications could be tested using a model in the following form:

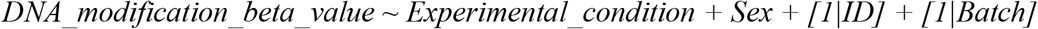

The modeling approach outlined above should be applicable to all types of paired 5-hmC and 5-mC data, provided the data structure is quantitative and at base-pair resolution. In addition, this approach should be appropriate for various analysis methods, provided they allow for a mixed effects design. Previous work has shown that beta regression (BR) and ratio of correlated gammas (RCG) modeling approaches are appropriate for detecting methylation differences on a genome-wide scale, and have greater specificity than linear models fitted to raw or normalized beta values (Triche et al., 2016; Weinhold et al., 2016). As such, fitting these types of models according to the repeated measures design outlined above should allow for simultaneous analysis of paired 5-mC and 5-hmC data, despite their potential differences in beta value distributions. Inclusion of an interaction term in the proposed model captures the potential transition from 5-mC to 5-hmC, allowing researchers to investigate whether experimental variables have distinct effects on 5-mC and 5-hmC dynamics at specific CpG sites/regions in neuronal tissue (see example in Figure 5). However, inclusion of an interaction term complicates interpretation of the main effect of experimental condition on the outcome of interest (i.e. DNA modification beta value). As a result, mixed effects models with an individual term for experimental condition, but no interaction term, can be used to model the response of either 5-mC or 5-hmC to experimental treatment. As the number of individual CpGs being tested increases, researchers must also consider instituting corrections for multiple testing – e.g. Benjamini-Hochberg false discovery rate (Benjamini and Hochberg, 1995). Future bioinformatics tools that aim to co-analyze paired 5-mC and 5-hmC data should implement this type of statistical approach on a genome-wide scale. This is particularly critical for epigenetics studies in brain tissue, where 5-hmC is both abundant and functionally relevant.

**Figure 5.**
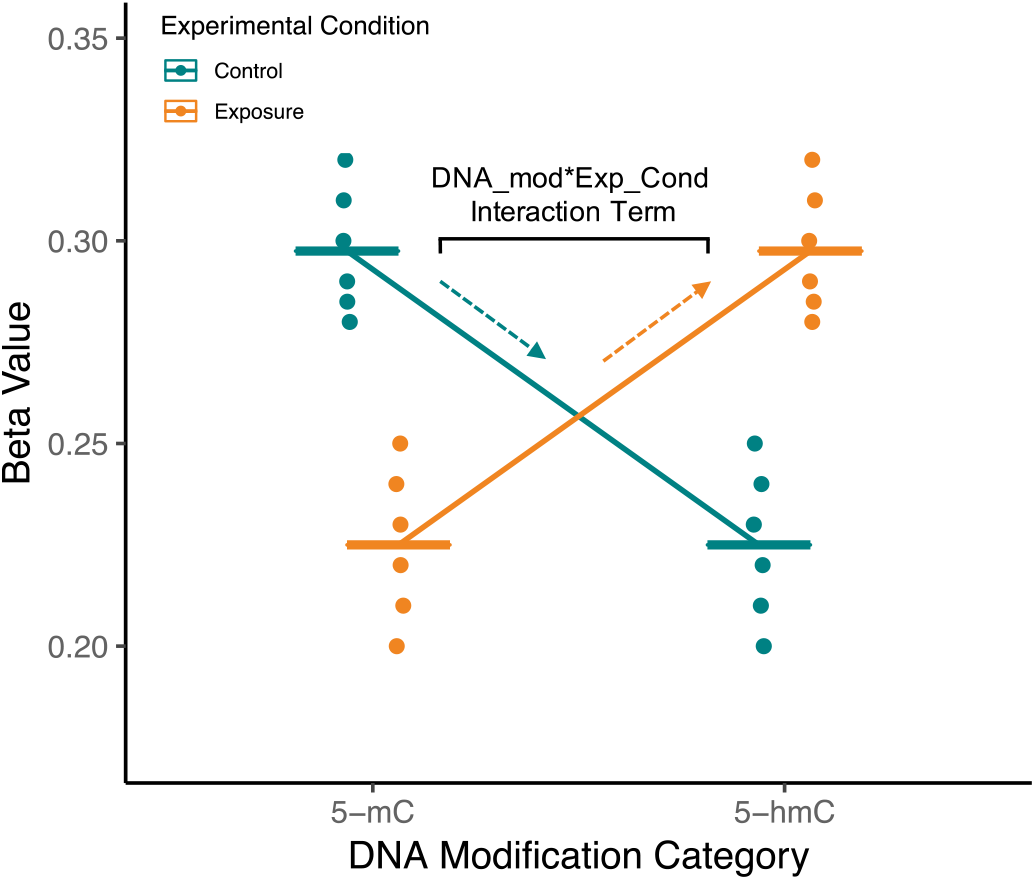
Visualization of interaction term from example repeated measures model for single CpG site from mock BS/oxBS data. In the proposed mixed effects model treating 5-mC and 5-hmC as repeated measures, a random effect for ID will account for the correlation between 5-mC and 5-hmC at a CpG site. Meanwhile, a DNA_mod*Exp_Cond interaction term will be used to determine whether 5-mC and 5-hmC differ in their response to experimental condition. In the visualized mock data, brain samples from exposed animals have an increased slope compared to control animals, indicating that exposure is shifting the CpG site toward 5-hmC in the brain. As indicated in the figure, this difference in slope is modeled by the DNA_mod*Exp_Cond interaction term. The proposed statistical approach can pick up regions of active DNA modification cycling while also accounting for the fact that 5-mC and 5-hmC are dependent measures. Furthermore, fixed effect terms for 5-hmC and 5-mC could also be included to model the response of either 5-mC or 5-hmC to experimental treatment.

## 5. Conclusion

Recent research has developed a number of methods for measuring genome-wide 5-hmC. These methods continue to improve and provide exciting new opportunities for understanding the biological role of DNA modifications. However, despite an abundance of available technical methods, it remains unclear how to best reconcile paired, base-pair resolution 5-mC and 5-hmC data. Here, we propose a statistical approach to handle 5-mC and 5-hmC as repeated measures using mixed effects models with an interaction term between experimental condition and DNA modification category. Using this approach, researchers would be able to investigate whether experimental variables have distinct effects on base-pair resolution 5-mC and 5-hmC in any collected tissue, including the brain. In this way, our proposed statistical method would allow for a deeper understanding of the interplay between 5-mC and 5-hmC in nervous system tissue, a necessary step on the road to designing targeted epigenetic therapeutics for neurological diseases.

## Notes

#### Summary of Updates

Corrected inaccurate statement in the abstract about the proposed modeling approach. The word linear was not accurate.

